## Supplementary Material for "Myelin dystrophy in the aging prefrontal cortex leads to impaired signal transmission and working memory decline: a multiscale computational study"

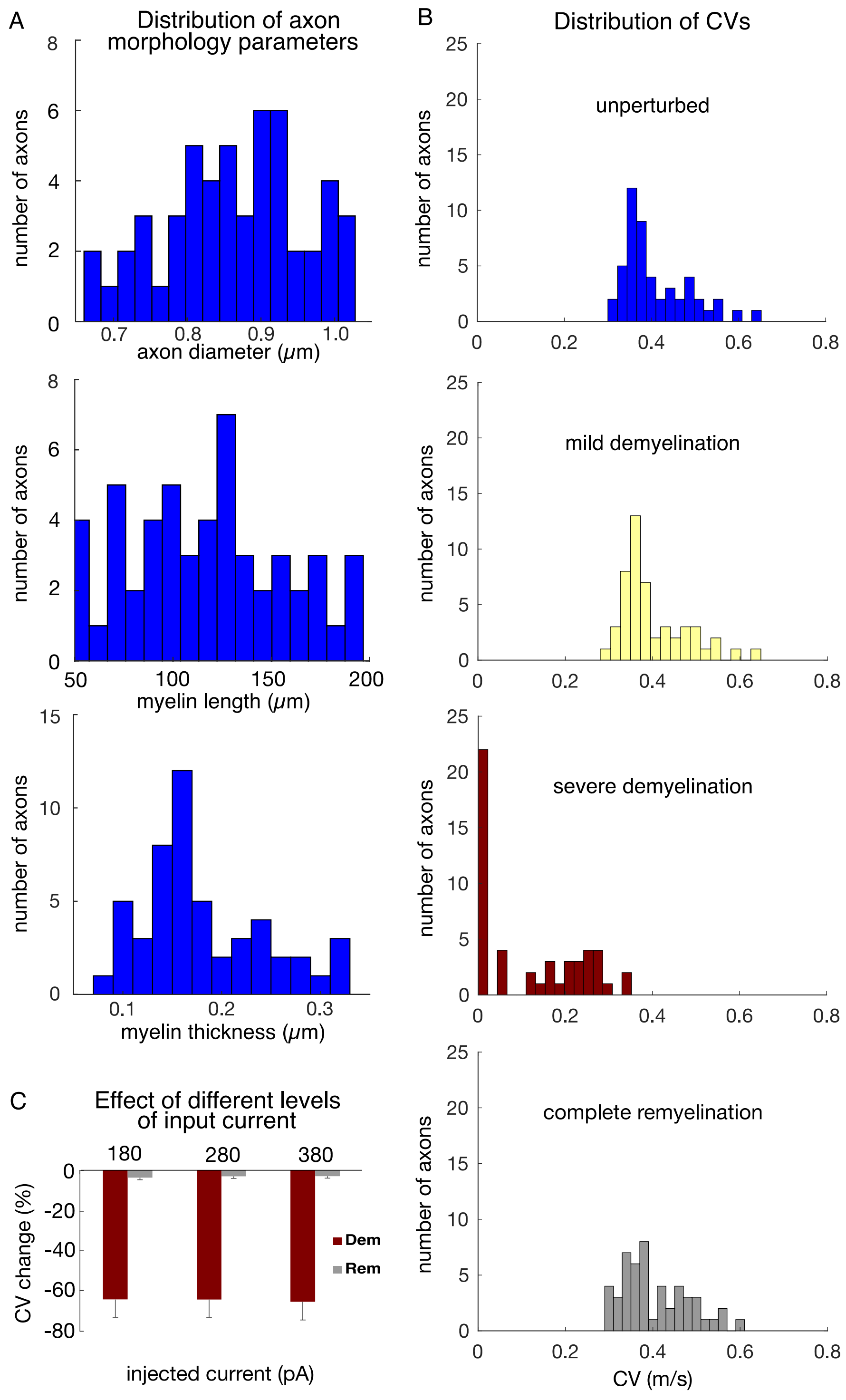


**Supplementary Figure 1. Distribution of parameters and conduction velocities in the single neuron model cohort.** (**A**) Histograms of axon morphology parameters of models selected for the single neuron cohort. Top: axon diameter; middle: length of unperturbed myelin segments; bottom: total myelin thickness in unperturbed segments, computed as the product of lamella thickness and number of lamellae. (**B**) Histograms of the CV for the 50 axons of the unperturbed model cohort (top), and representative demyelination and remyelination perturbations: mild demyelination (removing 25% of lamellae from 25% of the myelinated segments, second row); severe demyelination (removing all lamellae from 75% of the myelinated segments, third row); and complete (100%) remyelination (where the demyelinated segments from the third row were remyelinated by two shorter segments with 75% of lamellae). CVs averaged over 30 trials in each case. (**C**) Changes in CV (measured in %) in response to demyelination and remyelination versus the magnitude of current clamp step (+180, +280, or +380 pA). Shown are mean $\pm$ SEM for demyelinating 50% of myelinated segments (removing all lamellae), and subsequent remyelination of those segments by shorter segments with 75% of lamellae.

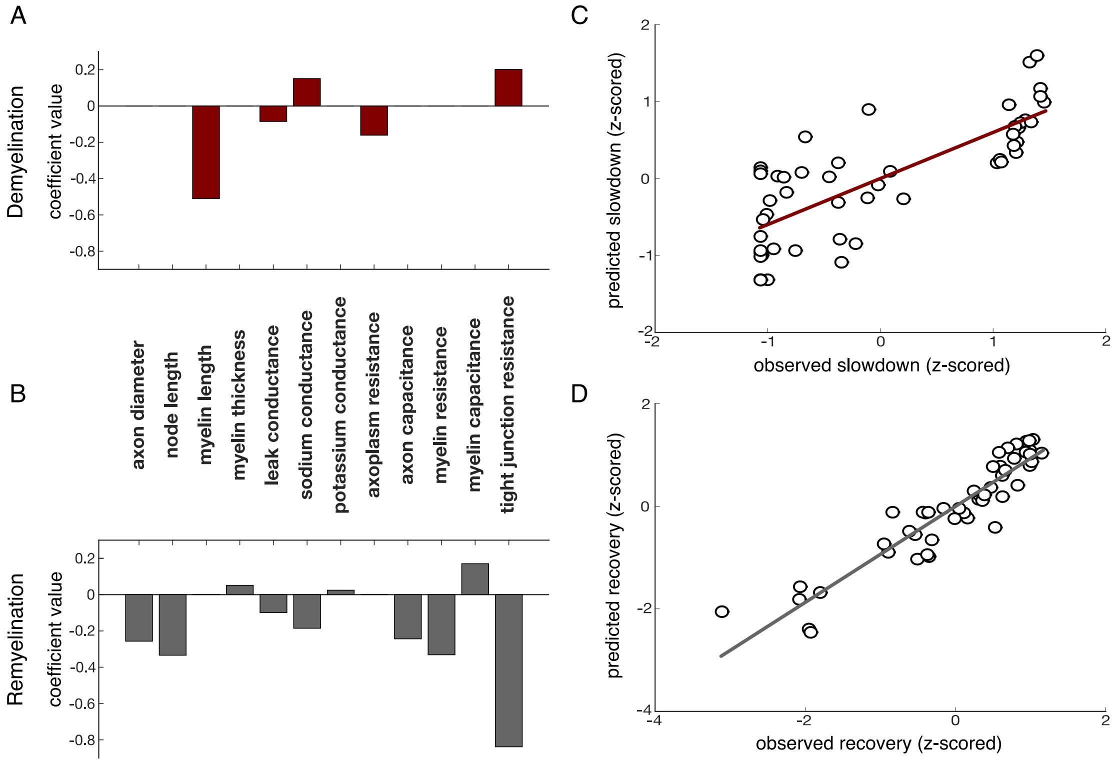


**Supplementary Figure 2. Statistical analysis of parameters contributing to CV changes after demyelination and remyelination.** Coefficients of Lasso regression models with 10-fold cross validation for demyelination (**A**) and remyelination (**B**). Parameters with non-zero coefficients are important factors underlying the response, critical in ascertaining the susceptibility of axons to respective perturbations. (**C-D**) The Lasso models effectively predicted how demyelination and remyelination affect CV. The models from **A**-**B** were applied to a novel test set (50 axons). Shown are predicted versus observed CV changes (z-scored; slowdown due to demyelination in **C**, recovery due to remyelination in **D**) for the 50 novel axons. Adjusted $R^2$ = 0.61 (**C**) and 0.87 (**D**) respectively.

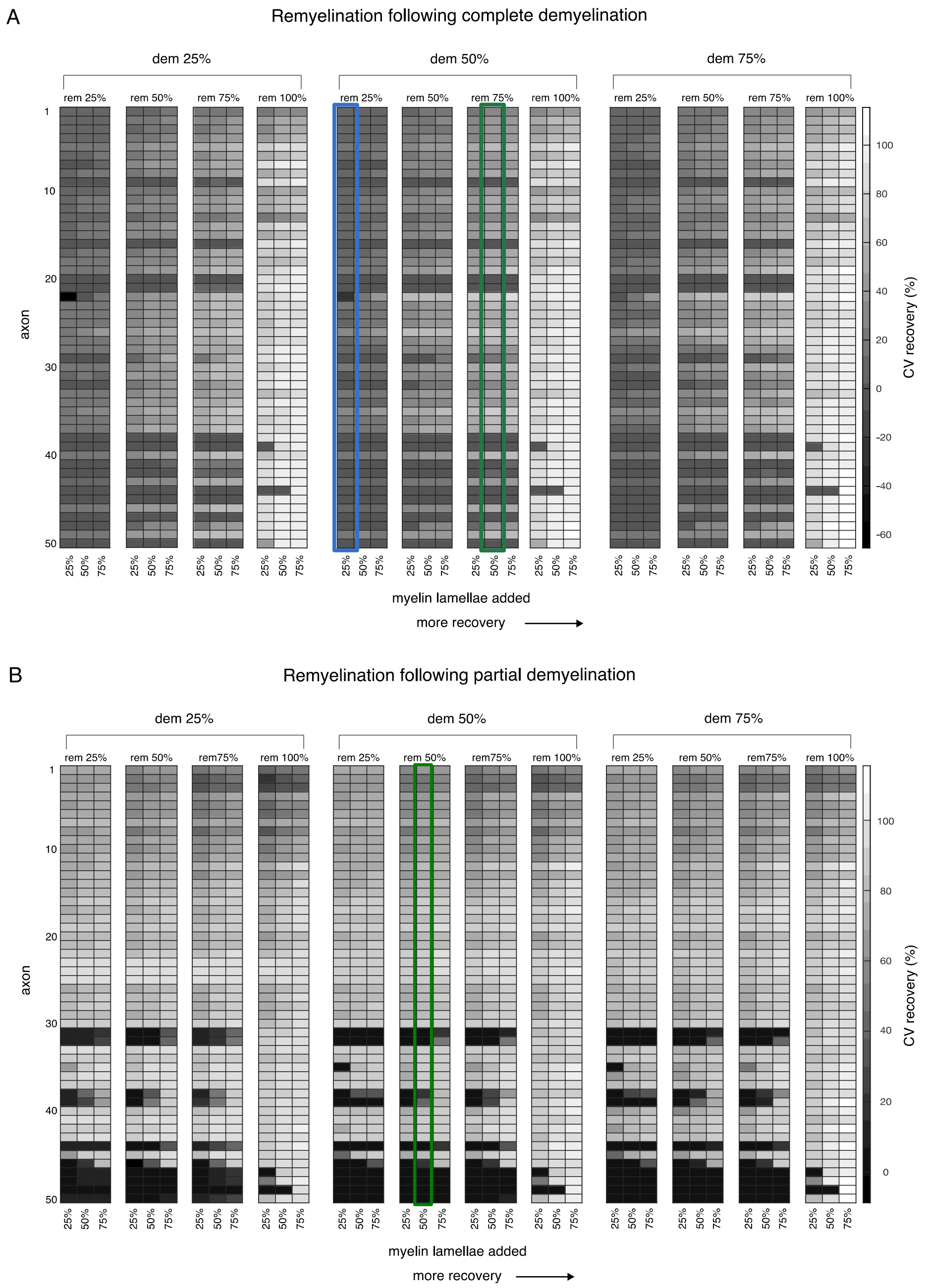


**Supplementary Figure 3. CV recovery in response to remyelination across the model cohort.** Axons arranged vertically in increasing order of myelinated segment length (longest at the bottom). The 3 groups of heat maps from left to right represent how many segments were demyelinated initially (25%, 50%, and 75% respectively). Within each group, the position in each of the four blocks indicates what proportion of demyelinated segments were remyelinated. Individual columns correspond to the percentage of lamellae restored to each remyelinated segment. Color of each box indicates the mean CV recovery from the corresponding demyelinated case across 30 trials (see Methods). **(A)** Recovery after complete demyelination, when all lamellae had been removed from affected segments. Columns highlighted in blue and green respectively correspond to the two remyelination cases shown in **Figure 4A**. CV recovery was positive, representing improvement, in all remyelination cases except one. For axon 22, the CV was worse for the mildest remyelination situation: when 25% of segments were initially demyelinated, then 25% of those segments were remyelinated by restoring just 25% of the myelin lamellae. A few axons showed recovery *above* 100%, suggesting faster conduction than in the unperturbed condition, when either 75% of the lamellae were restored. (**B**) Recovery after partial demyelination, when half of lamellae had been removed from affected segments. Column highlighted in green corresponds to the remyelination case shown in **Figure 4D.** CV recovery was positive, representing improvement, in all remyelination cases except one (axon 46, far left simulation condition).

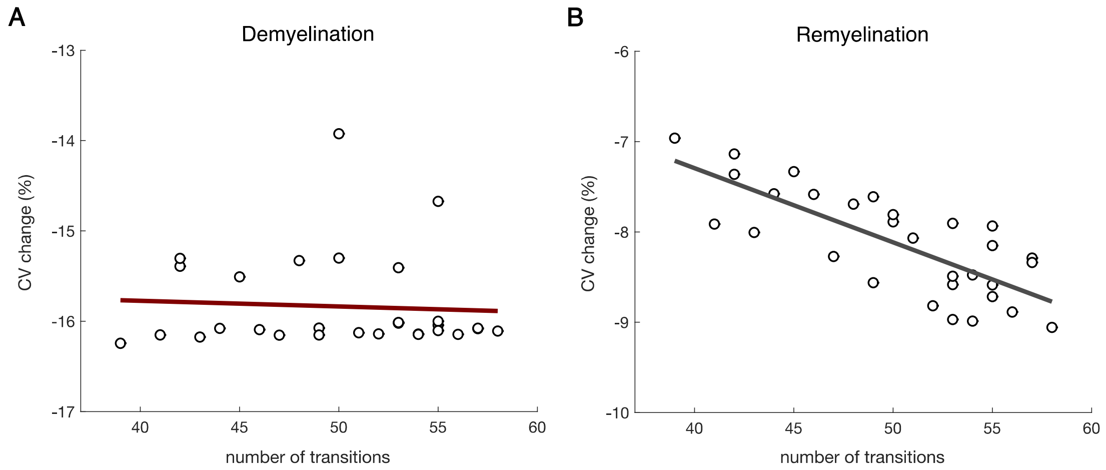


**Supplementary Figure 4. Transitions between myelinated segments of dissimilar lengths affect response to perturbations.** (**A**) Mean CV change (measured in %) versus the number of transitions from unperturbed to demyelinated segments in 30 randomized trials for a given demyelination condition (50% of the segments affected, 50% of lamellae removed). (**B**) Mean CV change (measured in %) versus the number of transitions from unperturbed (long) to remyelinated (short) segments in 30 randomized trials for complete remyelination of 50% of the segments with 50% of lamellae restored. CV change was more severe as the number of transitions between segments of unequal lengths increased.

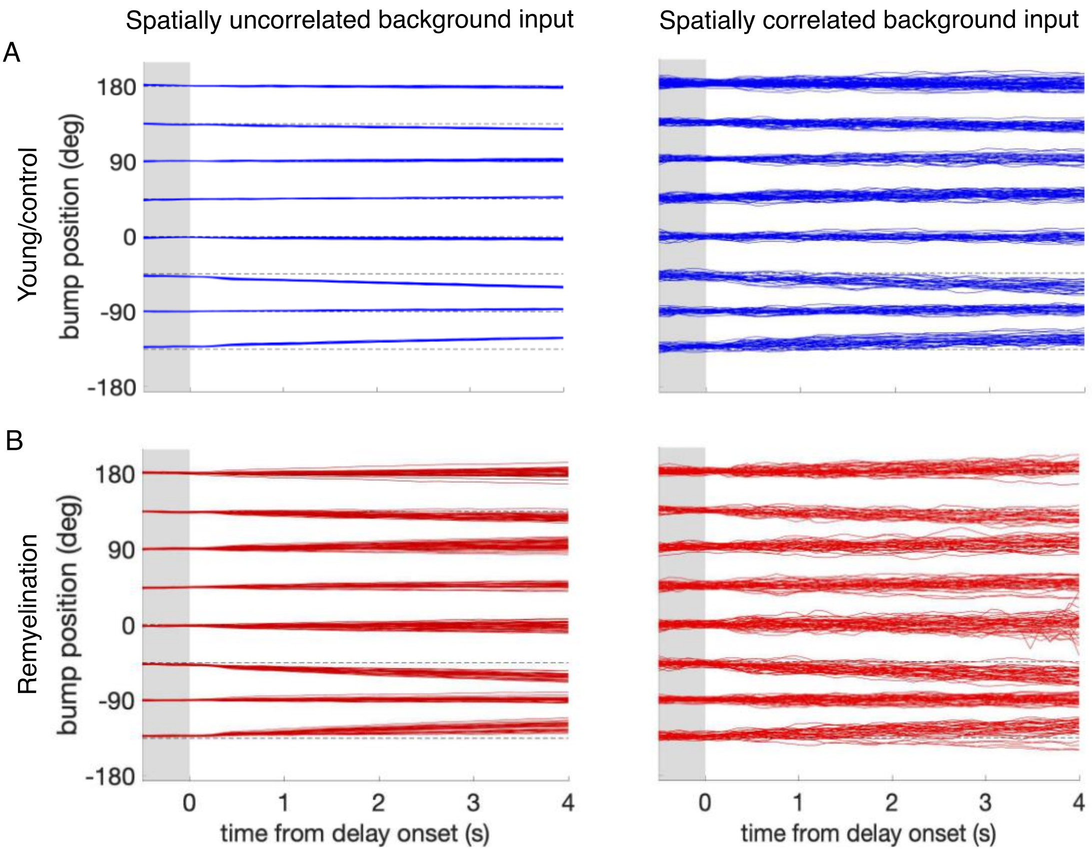


**Supplementary Figure 5. Increased working memory diffusion in spiking networks with spatially correlated background inputs.** Bump position during the cue and delay periods for 280 trials and the eight possible cue directions for (**A**) a young, control network and (**B**) a perturbed network when remyelinating 50% of the segments after partial demyelination of 25% of the segments along the neuronal axons, by adding 75% of the myelin lamellae back. Simulations were done with spatially uncorrelated background inputs as in all other simulations in **Figures 5-8** (left panels), and with spatially correlated background inputs (see Supplementary Methods; right panels). These simulations show that, as expected from theory, bump diffusion increases for spatially-modulated noise correlations (**Rosenbaum et al., 2017**; **Stein et al., 2021**). Importantly, the effect of myelin alterations is an increase in diffusion in both cases, showing the robustness of our results (diffusion constant = 0.018 deg$^2$/s in (**A**) left panel; 0.957 deg$^2$/s in (**B**) left panel; 3.115 deg$^2$/s in (**A**) right panel; and 9.605 deg$^2$/s in (**B**) right panel).

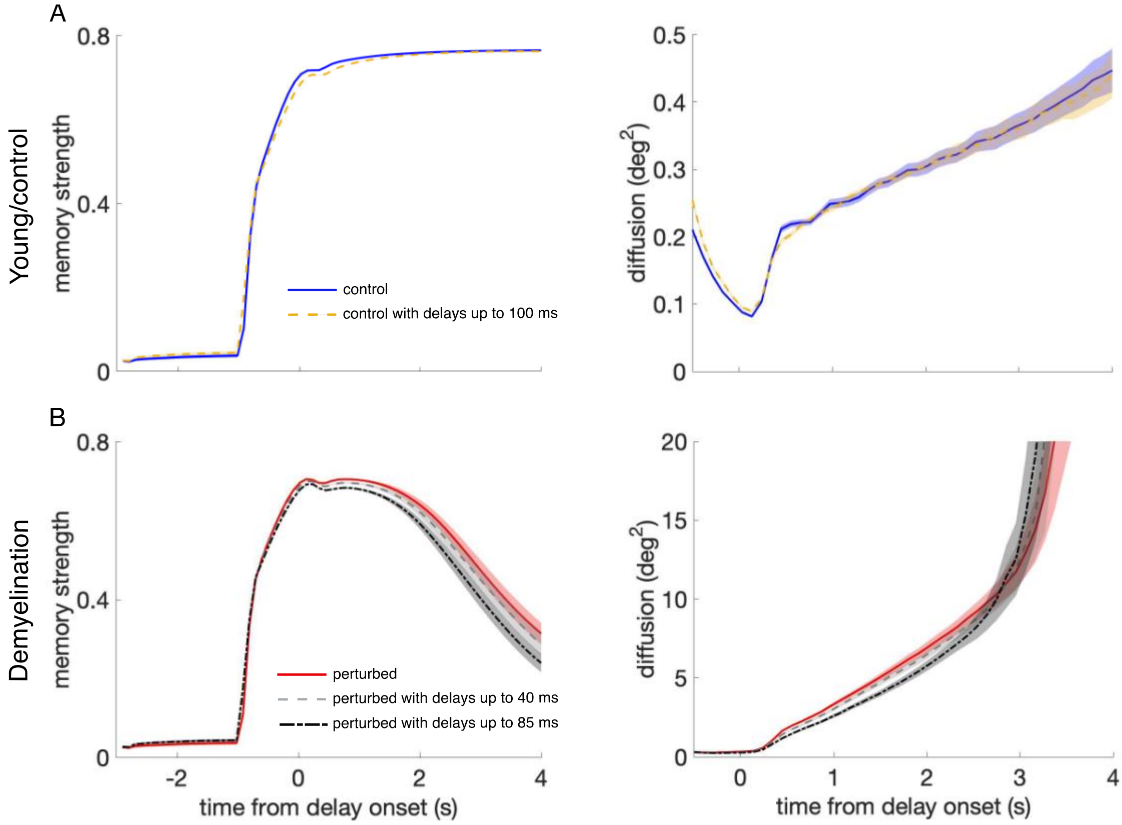


**Supplementary Figure 6**. **Effect of propagation delays on control and perturbed networks.** (**A**) Memory strength (left panels) and diffusion (right panels) for the young, control networks with zero propagation delays (blue solid line), as in **Figure 5**, and with propagation delays from a uniform distribution with a range between 0 and 100 ms (yellow dashed line). (**B**) Memory strength and diffusion for perturbed networks when demyelinating 50% of the segments along the axons of model neurons, by removing 60% of the myelin lamellae without delays (red solid line), and with delays from a uniform distribution with a range between 0 and 40 ms (gray dashed line) and between 0 and 85 ms (black dash-dotted line). The measures of working memory performance were calculated by averaging across 20 networks and 280 trials for each network. Shaded areas indicate SEM for each case. For the young, control networks, there was no difference with and without propagation delays, even though the delays used in the network simulations were much larger than the delays quantified in the single neuron model (the longest delays found for the most extreme perturbation condition – demyelination of 75% of the segments by removing 100% of the myelin lamellae– were of 49.9 ms on average; **A**). Working memory performance was also unaffected in the perturbed network with AP failures for delays ranging between 0 and 40 ms, also larger than the ones quantified in the single neuron model (for the case of 50% of the segments demyelinated by removing 60% of the myelin lamellae, the average delay in the cohort was 4.6 ms and the maximum delay was 15.7 ms; **B**). However, including extremely long delays of up to 85 ms did further impair memory compared to the impairment level introduced by AP failures alone (**B**).

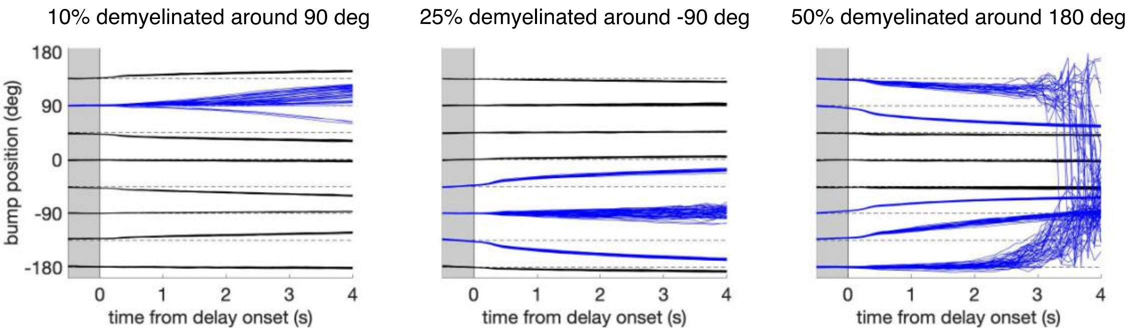


**Supplementary Figure 7. Effect of spatially heterogeneous demyelination of the model neurons according to their preferred angle.** We also tested working memory performance in the network when demyelination affects only parts of the network. The figure shows the decoded bump center position during the cue and delay period for the eight possible cue directions when a fraction of neurons was perturbed and the rest of the neurons in the circuit were unaltered (**Figure 5B**). We perturbed 10% of the neurons around the neuron with preferred direction 90° (left panel), 25% of the neurons around -90° (middle panel), and 50% of the neurons around 180° (right panel). Bump traces for cues that lie inside the perturbed portion of the circuit are shown in blue. Network perturbation in the three cases consisted in demyelinating 25% of the segments along the axons of model neurons, by removing 70% of the myelin lamellae. In each case, 280 trials were simulated for one network. These simulations show an increased drift and diffusion inside the perturbed zone, consistent with the increased drift and diffusion when perturbing the entire network (**Figure 6B** and **Supplementary Figure 11**). In particular, spatially heterogeneous demyelination in our network leads to a bias away from the affected zone and to increased trial-to-trial variability. Note that this is a model prediction, but we are not aware of empirical data showing heterogeneous demyelination with aging. Further, note that while our network model has a topological ring structure, neurons in PFC are not anatomically arranged depending on their preferred features. Thus, spatially heterogeneous demyelination would likely affect neurons with different feature preferences (i.e., neurons throughout our ring model).

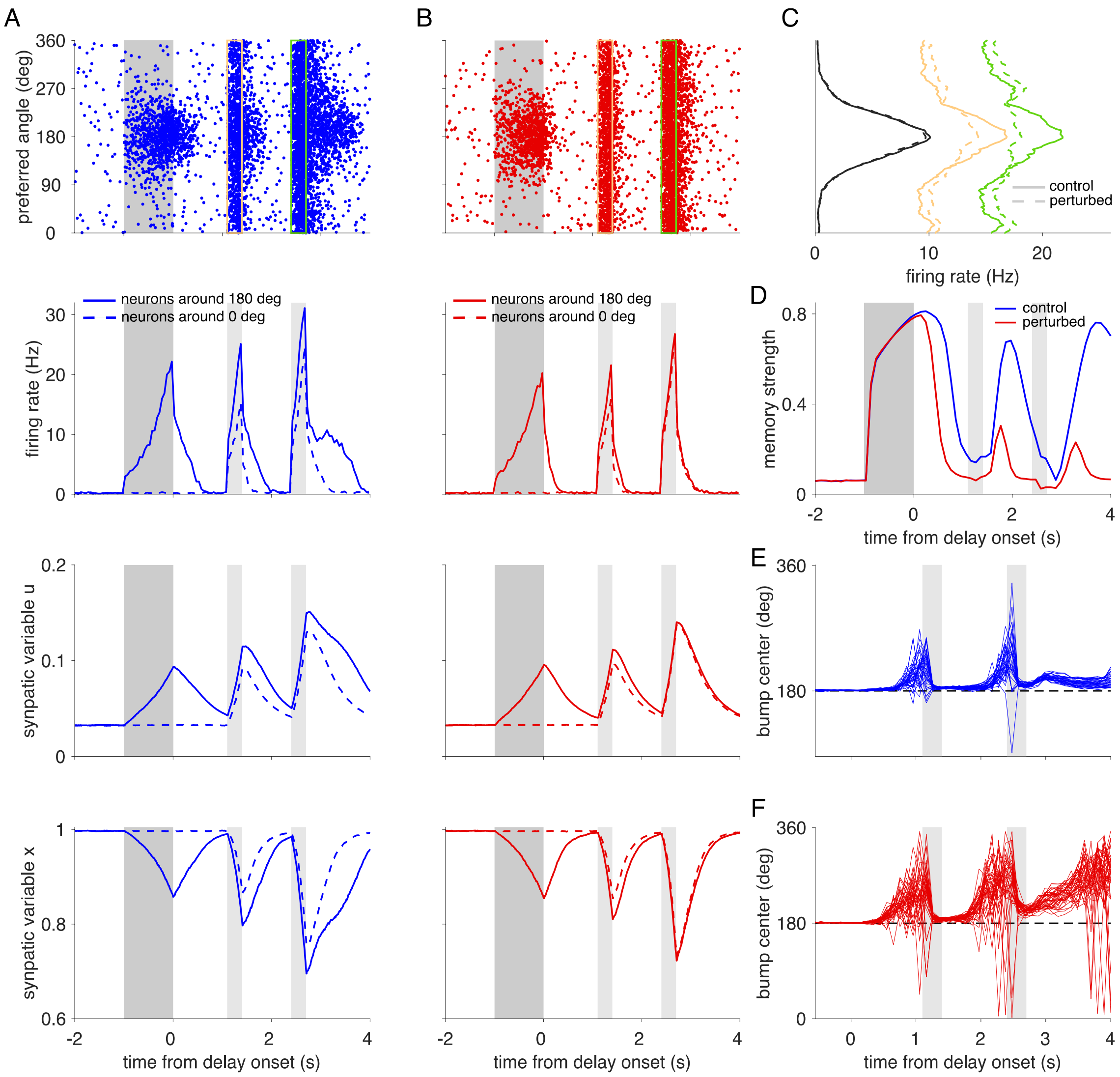


**Supplementary Figure 8. Action potential failures impair working memory performance in a network model with activity-silent memory traces.** (**A**) Spiking and synaptic activity in an unperturbed, activity-silent working memory model. Top: Raster plot showing the activity for each excitatory neuron (labeled by its preferred direction) in a single trial with a cue stimulus presented at 180°. We modified our spiking neural network model such that it does not show elevated persistent firing throughout the delay period (see **Figure 5B** for comparison). In particular, we reduced the external background input to excitatory neurons $I_E^{ext}$ by a factor of 3.61% and we increased the cue stimulus amplitude by 12.5%. Even though spiking activity decays to baseline (close to 0 Hz), a memory trace is imprinted in enhanced synaptic strength due to short-term synaptic facilitation (**Mongillo et al., 2008**). Selective spiking activity is recovered by a non-selective constant input applied during 300 ms to all excitatory neurons during the two reactivation periods (marked by yellow and green rectangles in the raster plot). The amplitude of the input was 11 mV during the first and 13 mV during the second reactivation period. Reactivation periods are marked in light gray shading in the remaining panels below and the cue period is indicated by dark gray shading. Firing rates (second row), synaptic facilitation variable u (third row), and synaptic depression variable x (bottom row) for the same trial, averaged for 500 neurons around the neuron with 180° as preferred direction (solid lines) and around the neuron with 0° as preferred direction (dashed lines). Note that reactivation recovers the activity bump (**C**) but also causes elevated firing and subsequent enhancement of synapses at all positions in the networks. (**B**) Activity in a network with demyelination of 50% of the myelinated segments by removing 60% of the myelin lamellae. AP failures lead to reduced firing rates in the cue and early delay periods and consequently to weaker synaptic enhancement. (**C**) Average spike counts of the excitatory neurons during the cue period (black lines), and the two reactivation periods indicated in the raster plots in **A** and **B** (yellow and green lines). Solid lines correspond to the control network and dashed lines to the perturbed network. (**D**) Memory strength as a function of time for the control and perturbed networks. (**E-F**) Trajectories of the bump center (i.e., remembered cue location) read out from the neural activity across the cue and delay periods using a population vector (see Methods). Cue position was 180° in all trials. The perturbed network (**F**) shows larger working memory errors towards the end of the delay period compared to the control network (**E**).

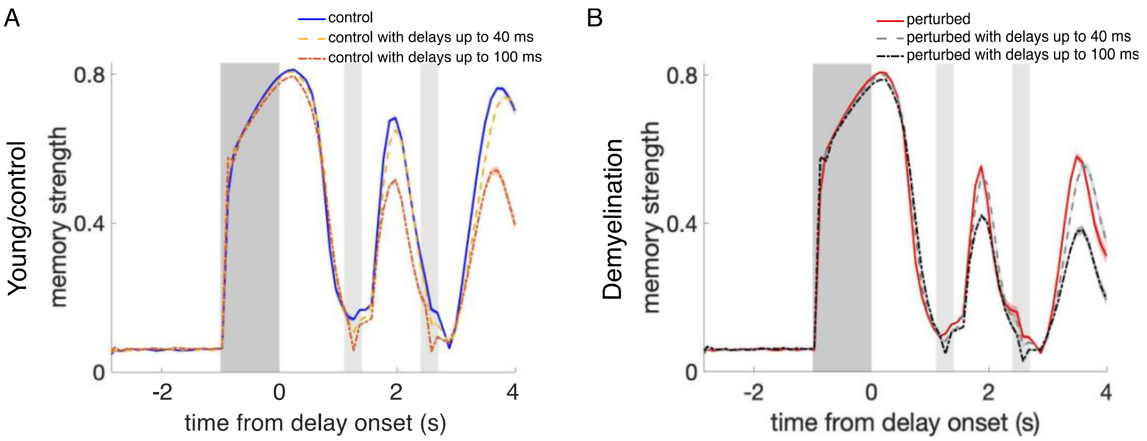


**Supplementary Figure 9. Effect of propagation delays on control and perturbed activity-silent network models.** (**A**) Memory strength during the whole simulation time for the young, control networks relying on activity-silent working memory (**Supplementary Figure 8**) with zero propagation delays (blue line), and with propagation delays from a uniform distribution with a range between 0 and 40 ms (yellow line) and between 0 and 100 ms (orange line). (**B**) Memory strength for perturbed networks when demyelinating 25% of the myelinated segments by removing 50% of the myelin lamellae, without delays (red line), and with uniformly distributed delays between 0 and 40 ms (light gray line) and between 0 and 100 ms (black line). The cue period is indicated by dark gray shading and reactivation periods are marked in light gray. Memory strength was calculated by averaging across 280 trials for one network. Shaded areas indicate SEM for each case. For the young, control networks (**A**), working memory was not affected by including delays of up to 40 ms. Unrealistically long delays ranging up to 100 ms did cause an impairment (the longest delays found for the most extreme perturbation condition – demyelination of 75% of the segments by removing 100% of the myelin lamellae – were of 49.9 ms on average). When also incorporating AP failures to the networks (**B**), we observed a similar trend. For this perturbation condition, delays of up to 40 ms were already much larger than the delays quantified in the single neuron model (for the case of 25% of the segments demyelinated by removing 50% of the myelin lamellae, the average delay in the cohort was 3.75 ms).

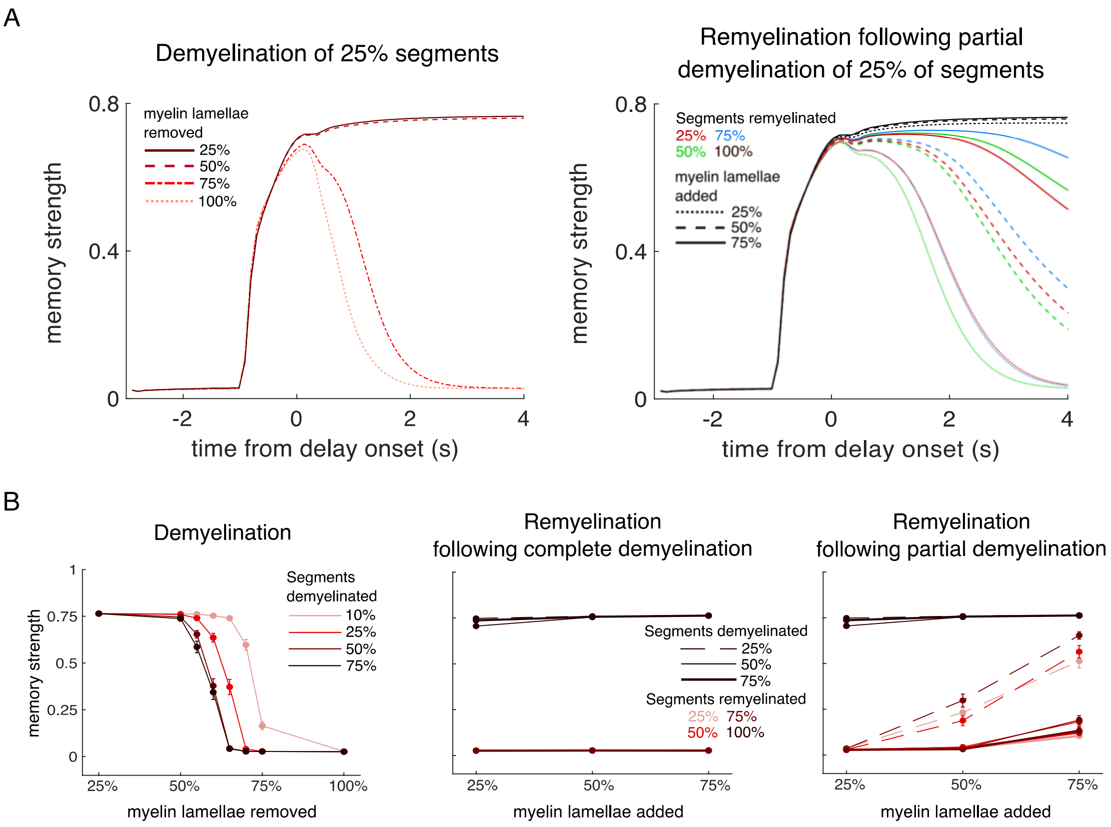


**Supplementary Figure 10. Memory strength decreases for different degrees of demyelination and remyelination.** (**A**) Progressive memory strength reduction, due to higher degrees of demyelination (left panel) and lower degrees of remyelination (right panel), leads to a progressive shortening of the memory duration. Left panel: demyelination of 25% of the myelinated segments by systematically removing myelin lamellae. Removing up to 50% of the lamellae causes no effect on the memory strength compared to the control networks (see **Figure 5C**; memory duration = 4 s). However, removing over 75% of the myelin lamellae progressively shortens memory duration (< 2 s). Right panel: systematic remyelination of the previously partially demyelinated 25% of the segments. Specially adding back more myelin lamellae recovers the memory strength and thus, the memory duration. Complete remyelination leads to control-like values (black lines; **Figure 5C**). (**B**) Memory strength at the end of the delay period for simulations of the DRT, for a systematic exploration of the effect of AP failure probabilities corresponding to the demyelination/remyelination conditions explored with the single neuron model (see **Figure 6**). Left panel: demyelination; middle panel: remyelination of the previously completely (removal of 100% of the myelin lamellae) demyelinated segments. Right panel: remyelination of the previously partially (removal of 50% of the myelin lamellae) demyelinated segments. In (**A**) and (**B**) memory strength was obtained by averaging across the 10 perturbed cohort networks and the 280 trials simulated for each network. The average memory strength for the 10 control networks in the cohort (averaged across 280 trials) was 0.765, corresponding to the case of 0% of lamellae removed in the left panel of (**B**). Error bars in (**B**) represent mean ± SEM, averaged across all networks and trials.

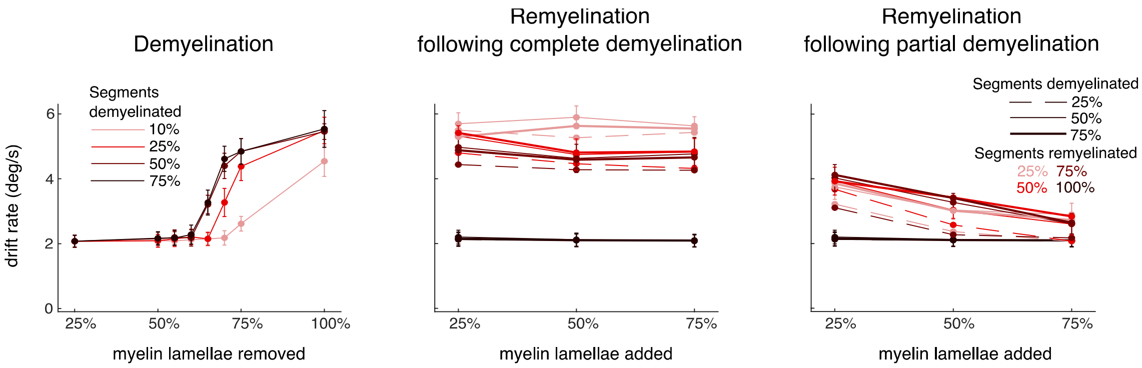


**Supplementary Figure 11**. **Increase of memory drift for different degrees of demyelination and remyelination.** Drift rate for simulations of the DRT, for a systematic exploration of the effect of AP failure probabilities corresponding to the demyelination/remyelination conditions explored with the single neuron model (see **Figure 6**). Left panel: demyelination; middle panel: remyelination of the previously completely (removal of 100% of the myelin lamellae) demyelinated segments. Right panel: remyelination of the previously partially (removal of 50% of the myelin lamellae) demyelinated segments. Drift rate was obtained by averaging across the 10 perturbed cohort networks and the 280 trials simulated for each network. The average drift rate for the 10 control networks in the cohort (averaged across 280 trials) was 2.075 deg/s, corresponding to the case of 0% of lamellae removed in the left panel. Error bars represent mean ± SEM, averaged across all networks and trials.

### Supplementary Methods

**Single cell model: Statistical assessment of parameter importance.**

To identify which parameters of the multicompartment model had the greatest influence on axonal responses to demyelination and remyelination perturbations, we used least absolute shrinkage and selection operator (Lasso) regression (**Tibshirani, 1996**; **James et al., 2021**) implemented in MATLAB R2022a (Mathworks, Natick, MA). Lasso fits a regression model to response variable, given predictor variables (observations of predictors) by selecting coefficients that minimize the quantity

$$\sum_{i=1}^{n}\left(y_i - \beta_0 - \sum_{j=1}^{p}\beta_j x_{ij}\right)^2 + \lambda\sum_{j=1}^{p}\left|\beta_j\right| \ ,$$

for a given value of (**James et al., 2021**). The former quantity is the residual sum of squares of the model. The latter quantity is the absolute sum of coefficients; including it in the minimization shrinks some coefficients to zero when is sufficiently large, enabling Lasso to perform feature selection. The tuning parameter was selected to minimize the 10-fold cross validation error; then for the selected value of, the regression was repeated using all available data. A total of 12 parameters served as predictor variables for the Lasso regression. We started with 6 of the 8 parameters used to construct the cohort (axon diameter, node length, myelin length, and scale factors for leak, NaF, and KDR conductances), and combined two others parameters (number of myelin lamellae and lamella thickness) into one quantity: myelin thickness (their product). Past studies (**Goldman and Albus, 1968**; **Koles and Rasminsky, 1972**; **Moore et al., 1978**; **Gow and Devaux, 2008**) identified several other parameters that affect axonal propagation, including the g-ratio, axoplasmic resistance, axon capacitance, myelin resistance, myelin capacitance, and tight junction resistance. We added five of these parameters to the seven cohort parameters as predictor variables, omitting only g-ratio due to its high correlation with axon diameter and myelin thickness. We created response variables summarizing the effects of demyelination and remyelination on CV in each of the 50 members of the model cohort. AP failures were recorded as CV change of -100%. We did not include AP failure rates as a response variable, since CV changes precede AP failures. For demyelination, we averaged the CV change for all randomized trials in which 100% of lamellae were removed from 25%, 50% and 75% of segments. For remyelination, we averaged the CV recovery in all randomized trials in which 25, 50, or 75% of segments were completely demyelinated, and then all affected segments were remyelinated as two shorter segments with 75% of lamellae added back. To facilitate comparison, we z-scored all predictor and response variables before performing Lasso separately on each response. We tested the predictive ability of the Lasso models by randomly selecting another 50 of the 138 models from the original hypercube that met the inclusion criteria, then simulating the specific myelin alteration protocols that comprised the demyelination and remyelination response variables. We computed Pearson's correlation coefficient between the z-scored predicted vs. observed responses.

**Network Model**

The following table summarizes all the parameters used in the network model simulations (**Figures 5-8**).

| Parameter | Value |
|---|---|
| $g_{Eea}$ | $533.3/\sqrt{K_E}\ mV \cdot ms$ |
| $g_{Een}$ | $490.64/\sqrt{K_E}\ mV \cdot ms$ |
| $g_{Eia}$ | $67.2/\sqrt{K_E}\ mV \cdot ms$ |
| $g_{Ein}$ | $7.4/\sqrt{K_E}\ mV \cdot ms$ |
| $g_{IE}$ | $-138.6/\sqrt{K_I}\ mV \cdot ms$ |
| $g_{II}$ | $-90.6/\sqrt{K_I}\ mV \cdot ms$ |
| $\tau_E$ | $20\ ms$ |
| $\tau_I$ | $10\ ms$ |
| $\tau_a$ | $3\ ms$ |
| $\tau_n$ | $50\ ms$ |
| $\tau_g$ | $4\ ms$ |
| $\tau_d$ | $200\ ms$ |
| $\tau_f$ | $450\ ms$ |
| $U$ | 0.03 |
| $V_T$ | $20\ mV$ |
| $V_R$ | $-3.33\ mV$ |
| $\sigma_{EE}$ | 30º |
| $\sigma_{EI}$ | 35º |
| $\sigma_{IE}$ | 30º |
| $\sigma_{II}$ | 30º |
| $I_E^{ext}$ | $1.66\ \sqrt{K_E}\ mV$ |
| $I_I^{ext}$ | $1.5355\ \sqrt{K_E}\ mV$ |
| $I_{max,E}$ | $0.24\ mV$ |
| $\varepsilon_E$ | 61.2º |

**Supplementary Table 1.** Network model parameters.

**Network simulations with spatially modulated correlations**

To introduce spatially modulated correlations in the model (**Supplementary Figure 5**), we reduced the strength of the constant background inputs to excitatory and inhibitory neurons, $I_E^{ext}$ and $I_I^{ext}$, by a factor of 0.5 and provide additional external input from a population of Poisson neurons. The parameters were chosen such that the time-averaged total input (the sum of the reduced constant input and the inputs from the Poisson population) is the same as in our default network without Poisson inputs. The external population is composed of $N^{ext}$ = 16,000 Poisson neurons that fire with a constant firing rate of $r^{ext}$ = 9.72 Hz. These neurons are connected through AMPA synapses to excitatory neurons in the network with a strength of $g_{Ea}^{ext} = 0.5\ g_{Eia}$ and to inhibitory neurons with a strength of $g_{Ia}^{ext} = 0.4625\ g_{Eia}$. The average total number of synaptic inputs from Poisson neurons that a neuron receives is $K^{ext}$ = 1,000. Crucially, the connections from the Poisson population to neurons in the network are spatially structured, with a Gaussian connection profile similar to the recurrent connections in the network (ring structure) and with the interaction range determined by $\sigma^{ext}$ = 20º. Thus, neurons in the network receive shared inputs with a spatial structure and this leads to spatial correlations in the network neurons (**Rosenbaum et al., 2017**).
